# Probabilistic atlas of the lateral parabrachial nucleus, medial parabrachial nucleus, vestibular nuclei complex and medullary viscero-sensory-motor nuclei complex in living humans from 7 Tesla MRI

**DOI:** 10.1101/814228

**Authors:** Kavita Singh, Iole Indovina, Jean C. Augustinack, Kimberly Nestor, María G. García-Gomar, Jeffrey P Staab, Marta Bianciardi

## Abstract

The lateral parabrachial nucleus, medial parabrachial nucleus, vestibular nuclei complex and medullary viscero-sensory-motor nuclei complex (the latter including among others the solitary nucleus, vagus nerve nucleus, and hypoglossal nucleus) are anatomically and functionally connected brainstem gray-matter structures that convey signals across multiple modalities between the brain and the spinal cord to regulate vital bodily functions. It is remarkably difficult to precisely extrapolate the location of these nuclei from *ex vivo* atlases to conventional 3 Tesla *in vivo* images; thus, a probabilistic brainstem atlas in stereotaxic neuroimaging space in living humans is needed. We delineated these nuclei using single-subject high contrast 1.1 mm isotropic resolution 7 Tesla MRI images. After precise coregistration of nuclei labels to stereotaxic space, we generated a probabilistic atlas of their anatomical locations. Finally, we validated the nuclei labels in the atlas by assessing their inter-rater agreement, consistency across subjects and volumes. We also performed a preliminary comparison of their location and microstructural properties to histologic sections of a postmortem human brainstem specimen. In future, the resulting probabilistic atlas of these brainstem nuclei in stereotaxic space may assist researchers and clinicians in evaluating autonomic, vestibular and viscero-sensory-motor nuclei structure, function and connectivity in living humans using conventional 3 Tesla MRI scanners.

## 1. Introduction

The lateral parabrachial (LPB) nucleus, medial parabrachial (MPB) nucleus, vestibular (Ve) nuclei complex and medullary viscero-sensory-motor (VSM) nuclei (i.e. solitary nucleus, vagus nerve nucleus, hypoglossal nucleus, prepositus, intercalated nucleus, and interpositus) complex are anatomically and functionally connected brainstem gray-matter structures that convey signals across multiple modalities between the brain and the spinal cord to regulate vital bodily functions. Specifically, these structures, individually or synergistically, regulate arousal (e.g. LBP, MPB) (Kaur et al. 2013), gustatory processes (e.g. VSM) (Matsumoto 2013), sensory-motor function (VSM) (Olszewski and Baxter 1954), and autonomic functions like cardio-respiratory (e.g. LPB, MPB) (Damasceno, Takakura, and Moreira 2014) and gastrointestinal processes (e.g. VSM) (Bokiniec et al. 2017). Clinical conditions that alter the structure or function of these nuclei, including cerebrovascular events, autoimmune diseases, trauma, stroke (Choi and Kim 2018)and neurodegenerative disorders may produce a wide variety of symptoms and signs including disruptions of sleep and alertness, autonomic dysregulation, vertigo, and impaired control of eye movements and gait.

To identify the location of arousal, vestibular and viscero-sensory motor nuclei in living humans, neuroscientists and neurosurgeons currently rely on the work of neuroanatomists and pathologists, who created meticulous *postmortem* atlases of the human brainstem (Olszewski and Baxter 1954; Paxinos and Huang 1995; Naidich et al. 2009; Paxinos et al. 2012). Yet, it is remarkably difficult to precisely extrapolate the location of these nuclei from *ex vivo* atlases to conventional 3 Tesla *in vivo* images because these nuclei are not clearly visible in conventional imaging and display inter-subject and age-dependent variability. Moreover, *postmortem* atlases (Olszewski and Baxter 1954; Paxinos and Huang 1995; Naidich et al. 2009; Paxinos et al. 2012) are mostly *2D, non-probabilistic* (i.e. derived from a single or very few brainstem specimens —mostly of elderly subjects), and *non-deformable* (i.e. mostly static) representations of brainstem nuclei with inherent distortions due to tissue manipulation.

This indicates the need to expand current probabilistic neuroimaging brain atlases (Desikan et al. 2006; Destrieux et al. 2010; Bianciardi et al. 2015, 2018), to include these currently missing nuclei relevant for the aforementioned broad set of diseases.

The aim of this study was to create, in living humans, a stereotaxic probabilistic structural atlas of the right and left LPB (LPB-r, LPB-l), MPB (MPB-r, MPB-l), Ve (Ve-r, Ve-l) and VSM (VSM-r, VSM-l) using high-resolution (1.1-mm isotropic), multi-contrast diffusion fractional-anisotropy (FA) and T_2_-weighted images at 7 Tesla. This tool may augment modern day research and clinical brainstem studies by enabling a more precise identification of the location of these nuclei in conventional 3 Tesla images of living humans.

## 2. Material and methods

### 2.1 MRI data acquisition

To delineate LPB, MPB, Ve, and VSM nuclei we used data acquired in our previous study (Bianciardi et al., 2015), which is detailed below. After providing written informed consent in accordance with the Declaration of Helsinki, twelve healthy subjects (6m/6f, age 28 ± 1 years) underwent 7 Tesla MRI. The Massachusetts General Hospital Institutional Review Board approved the study protocol. Data were acquired using a custom-built 32-channel receive coil and volume transmit coil (Keil et al. 2010), which provided increased sensitivity for the brainstem compared to commercial coils. Common single-shot 2D echo-planar imaging (EPI) scheme was utilized to obtain 1.1 mm isotropic diffusion-tensor images (DTI) in sagittal plane, and T_2_-weighted images, with following parameters: matrix size/GRAPPA factor/nominal echo-spacing = 180 × 240/3/0.82 ms. The resulting T_2_-weighted anatomical images and the DTI images had perfectly matched resolution and geometric distortions. The EPI scheme helped us overcome specific-absorption-rate limits of spin-warp T_2_-weighted MRI at 7 Tesla. The additional parameters used for DTI and T_2_-weighted image acquisition included: spin-echo EPI, echo-time/repetition-time = 60.8 ms/5.6 s, slices = 61, partial Fourier: 6/8, unipolar diffusion-weighting gradients (for DTI), number of diffusion directions = 60 (for DTI, b-value ~ 1000 s/mm^2^), 7 interspersed “b0” images (non-diffusion weighted, b-value ~ 0 s/mm2, also used as T_2_-weighted MRI), 4 repetitions, acquisition time per repetition 6’43”. The entire acquisition time for T_2_-weighted MRI and DTI was ~ 27’. Importantly, for DTI acquisition at 7 Tesla, where tissue has shorter T_2_ values, we used unipolar (Stejskal and Tanner 1965) instead of bipolar (Reese et al. 2003) diffusion gradients. This led to shortened echo-time (less by ~30 ms) and significantly improved sensitivity of high-resolution DTI.

### 2.2 MRI data pre-processing and alignment to MNI space

On a single subject basis, after concatenation of four DTI repetitions, the data were preprocessed for distortion and motion artifacts using the Diffusion Toolbox in the FMRIB Software Library (FSL, Oxford, UK). The diffusion tensor at each voxel was estimated (using FRIMB’s Diffusion toolbox) to compute diffusion tensor FA from tensor eigenvalues. After motion correction, affine transformation was performed to coregister the averaged 28 “b0” T_2_-weighted images to DTI data (Bianciardi et al., 2015).

On a single-subject basis, precise co-registration of both T_2_-weighted images and FA to MNI space was performed as in (Bianciardi et al. 2015). Specifically, the brainstem of each subject was aligned to an MNI-space based FA template (termed “IIT space”) (Illinois Institute of Technology human brain atlas, v.3, Chicago, USA) (Varentsova, Zhang, and Arfanakis 2014) using the Advanced Normalization Tool (ANTs, Philadelphia, USA) (Avants et al. 2011). This template was used since it has high contrast, is compatible with diffusion-based tractography and covers the whole brainstem (encompassing medulla as well). Particularly, we computed and concatenated a generic affine and a high-dimensional non-linear warp transformation of images with the same modality (FA maps). The generic affine transformation was calculated by concatenating center-of mass alignment (degrees of freedom-dof = 3), rigid (dof = 6), similarity (dof = 7) and fully affine (dof = 12) transformations with smoothing sigmas: 4, 2, 1, 0 voxels – fixed image space. The high-dimensional non-linear warp transformation employed histogram image matching prior to registration and a symmetric diffeomorphic normalization transformation model with smoothing sigmas: 3, 2, 1, 0 voxels – fixed image space. We also performed a cross correlation metric, gradient step size: 0.2; regular sampling, data winsorization – quantiles: 0.005, 0.995; four multi-resolution levels: shrink factors 6, 4, 2, 1 voxels – fixed image space; convergence criterion: slope of the normalized energy profile over the last 10 iterations < 10^−8^. The resulting combined transformation (using a single-interpolation step method: linear), was then applied to both single-subject T_2_-weighted and FA images. Further, for each subject, T_2_-weighted and FA images were also aligned to MNI152 standard space (non-linear 6th generation MNI152_T1_1mm available for instance in FSL) (coined “MNI152_1mm space”), which is a frequently utilized space for fMRI analysis. While the MNI152_1mm space and the IIT space show satisfactory alignment elsewhere, there is slight misalignment in the brainstem, particularly in the pons and medulla. As such, single-subject FA and T_2_-weighted images were aligned to MNI152_1mm space. This was done by using a single-interpolation step (interpolation method: linear) and applying two concatenated transformations, namely single-subject to IIT space transformation (described above); and IIT to MNI152_1mm non-linear transformation, with parameters described above.

### 2.3 Single-subject labeling and probabilistic atlas generation

On a single-subject basis, two raters (K.S. and M.B.) independently performed manual delineations (fslview, FSL, Oxford, UK) using multi-contrast FA maps and T_2_-weighted images in IIT space to yield single-subject labels (i.e. masks) of the regions of interest (LPB-r/l, MPB-r/l, Ve-r/l and VSM-r/l). The intersection of the labels of the two raters was used as the final label. Manual delineations were aided by the use of anatomical landmarks and neighborhood rules described in a literature postmortem brainstem atlas (Paxinos et al. 2012). Of note, to delineate each nucleus, we mainly used the image modality that displayed the nucleus boundaries with good contrast (FA MRI for LPB, MPB, Ve, VSM), and employed the other modality (T_2_-weighted), which had poor contrast for that nucleus, to identify cerebrospinal fluid (CSF) boundaries (as for LPB and Ve nuclei).

A probabilistic neuroimaging atlas in IIT space was formulated for each nucleus as an average probability map of the nucleus label encompassing all subjects (100 % overlap of nuclei labels across subjects, *n* = 12 was considered highest probability). After registering the individual subject labels to MNI152_1mm space, by applying the IIT to MNI152 transformations described above (nearest neighbors interpolation), a similar atlas was derived in MNI152_1mm. We developed the resulting atlas (in both IIT and MNI152 spaces), to facilitate extrapolation to diffusion and functional MRI modalities.

For each subject and label (coregistered to single-subject native space via an inverse of the method described in section 2.2) we also calculated the label volume in native space, yielding the mean (s.e.) volume for all subjects and compared these values to literature (Paxinos et al. 2012) volumes as described in section 2.4.

### 2.4 Atlas validation

The probabilistic nuclei atlas was validated by computing for each nucleus and subject: (i) the inter-rater agreement, as the modified Hausdorff distance between labels delineated by the two raters; (ii) the internal consistency across subjects of the final label, as the modified Hausdorff distance between each final label and the probabilistic atlas label (thresholded at 35%) generated by averaging the labels across the other 11 subjects (leave-one-out cross validation). For both the inter-rater agreement and the internal consistency, we calculated the modified Hausdorff distance (Dubuisson and Jain 1994) which is a measure of spatial overlap frequently used in neuroimaging (Fischl et al. 2008; Klein et al. 2009, 2010; Augustinack et al. 2013). For each label, the minimum distance of every point on one label from the other label was averaged across all points, resulting in two distance values. The maximum value of these two values was calculated and used as modified Hausdorff distance. For each nucleus, the modified Hausdorff distance of (i) and (ii) was then averaged across subjects.

For further (iii) probabilistic atlas validation, we computed the volume of the delineated nuclei (i.e. of the final labels, intersection of the labels generated by each rater) and compared them to precisely computed literature nuclei volumes from the Paxinos atlas (Paxinos et al. 2012). For nuclei volume calculation based on the Paxinos atlas, we acquired snapshots of brainstem plates ranging from −5 mm to +32 mm (Figures 8.12-8.49 of (Paxinos et al. 2012) using Adobe Acrobat Reader. Later each snapshot was converted to single-slice nifti images with a slice thickness of 1 mm and proper spatial resolution using Matlab. To determine the in-plane isotropic spatial resolution of each slice, the number of pixels between adjacent coordinates was computed manually based on coordinate system provided for each plate (Paxinos et al. 2012). This varied between .0222 mm and .0417 mm in the examined plates. Based on Paxinos nomenclature (Paxinos et al., 2012), we manually delineated using ITK-Snap (Yushkevich et al. 2006) sub-regions of LPB, MPB, Ve and VSM and combined them to obtain final nuclei. For MPB, based on Paxinos terminology, we combined MPB and MPB external part (MPBE). For LPB, based on Paxinos atlas, we combined sub-nuclei of LPB external (LPBE), LPB central (LPBC), LPB dorsal (LPBD), LPB, LPB-superior (LPBS), and LPB unlabeled (i.e. a very small neighboring region compatible with LPB, yet missing the LPB label in the Paxinos atlas). We delineated and combined to final Ve nuclei complex the following Paxinos sub-nuclei: the nucleus of origin of vestibular efferents of the vestibular nerve (EVe), lateral vestibular nucleus (LVe), medial vestibular nucleus magnocellular part (MVeMC), medial vestibular nucleus (MVe), medial vestibular nucleus parvicellular part (MVePC), paravestibular nucleus (PaVe), spinal (i.e. inferior) vestibular nucleus (SpVe) and superior vestibular nucleus (SuVe). Similarly for VSM, we delineated and combined labels of: solitary nucleus commissural part (SolC), solitary nucleus dorsolateral part (SolDL), solitary nucleus dorsal part (SolD), solitary nucleus gelatinous part (SolG), solitary nucleus intermediate part (SolIM), solitary nucleus interstitial part (SolI), solitary nucleus medial part (SolM), solitary nucleus paracommissural part (SolPaC), solitary nucleus ventrolateral part (SolVL), solitary nucleus ventral part (SolV), vagus nerve nucleus (10N), hypoglossal nucleus (12N), prepositus (Pr), intercalated nucleus (In), and interpositus nucleus (IPo). To get a literature nucleus/sub-nucleus volume, we multiplied the number of delineated voxels in each nucleus/sub-nucleus by the voxel volume for each slice and added the resulting number across slices for each nucleus/subnucleus.

For additional (iv) validation of LPB and MPB probabilistic atlas labels, we performed preliminary histological evaluation of these nuclei in a postmortem human brainstem specimen. A brainstem specimen from a 65-year-old adult male without neurologic disease was obtained from MGH Autopsy Suite and studied. Five mesopontine transverse vibratome sections (50 μm thick) were Nissl (cell body) stained, and five adjacent sections were Gallyas (myelin) stained. Each section was mounted onto a gelatin dipped glass slide and dried overnight. <u>*Nissl thionin-based staining:*</u> After defatting with chloroform-alcohol, and undergoing pre-treatment with acetic acid, sections were stained with 1 % thionin for 3 minutes, differentiated in 70% with a few drops of glacial acetic acid, and dehydrated in ascending series (70 %, 70 %, 95 %, 95 %, 95 %, 100 %, 100 %) of alcohol. *Gallyas staining:* Sections were post-fixed with 10% formol for 10 days. Then, they underwent the following steps: 1) *acetylation* with 2:1 mixture of pyridine and acetic anhydride (30 minutes); 2) *wash* with distilled water; 3) *impregnation* with ammoniacal silver nitrate (pH 7.3, 30 minutes); 4) *wash* with distilled water; 5) *development* with stock ABC solutions of ammonium nitrate, sliver nitrate and tungstosilicic acid (10 minutes); 6) *development stop* with 1 % acetic acid; 7) *bleaching* with 0.2 % potassium ferricyanide (10 minutes); 8) *stop* with 0.5 % acetic acid; 9) *stop* with 0.5 % sodium thiosulfate; 10) *wash* with distilled water; 11) *dehydration* with ascending series (50 %, 70 %, 95 %, 100 %) of alcohol. <u>*Cover-slipping and digitization:*</u> After Nissl or Gallyas staining, sections were cover-slipped from xylene using Permount. The Nissl and Gallyas stained sections were digitized using an 80i Nikon Microscope (Microvideo Instruments, Avon, MA, USA) with a 4× objective (i.e. 40× total magnification), which resulted in images with a 1.85 μm pixel size. We automatically acquired the images using the virtual tissue workflow provided from Stereo Investigator (MBF Bioscience, Burlington, VT, USA). Later we subtracted the background using GIMP (https://www.gimp.org), an open-source drawing and annotation software.

## 3. Results

The probabilistic neuroimaging structural labels in MNI space of LPB-r/l, MPB-r/l, Ve-r//l and VSM-r/l are shown in Figures 1-4. We briefly describe the nuclei delineations on the basis of the MRI contrast that guided them, as well as on the basis of neighborhood relationships with other visible structures. LPB (Figure 1) appeared as a thin hypointense region on FA maps at the mesopontine junction, bounded medially and ventro-medially by the superior cerebellar peduncle (a hyperintense region in FA maps), rostrally by the pedunculotegmental nucleus (a hypointense region in FA maps described in Bianciardi et al 2018), and dorsally/dorsolaterally by the CSF (visible in T_2_-weighted MRI). On FA maps, the MPB (Figure 2) was visible as a thin hypointense stripe lying along the medial surface of the superior cerebellar peduncle caudal to its decussation in the lateral part of the oral pontine tegmentum, and extending down to the level of oral pole of superior vestibular nucleus (described below). The superior, medial, lateral and spinal vestibular nuclei were not visible as individual nuclei, yet they appeared as a single hypointense oblong-shaped region on FA maps, which was labeled as Ve nuclei complex (Figure 3). This complex extended from the caudal tip of the MPB at the ponto-medullary junction, to the medulla at the level of the mid inferior olivary nucleus (a hypointense region in FA, delineated in our previous study (Bianciardi et al. 2015)). On axial T_2_-weighted images, this complex was bounded dorso-laterally by the CSF of the fourth ventricle (visible in T_2_-weighted MRI). Finally, we delineated solitary along with nuclei 12N, 10N and smaller nuclei Pr, In, IPo within the VSM nuclei complex (Figure 4), an area of diffusion FA brighter than the adjacent medullary reticular formation and darker than neighboring white matter fiber bundles (e.g. medial longitudinal fasciculus). The VSM nuclei complex lied inferiorly to the ponto-medullary junction extending caudally throughout the medulla along the whole extent of the inferior olivary nucleus (hypointense region on FA images delineated in (Bianciardi et al. 2015)). On a coronal view, both VSM bilateral nuclei appeared as an inverted V-shape dorsally at their apex (see Figure 4). Further, VSM assumed a lateral position in the floor of fourth ventricle in the periventricular medullary region. Orally, it lied next to Ve as a hypointense region, as seen in axial views of FA maps.

**Figure 1.**
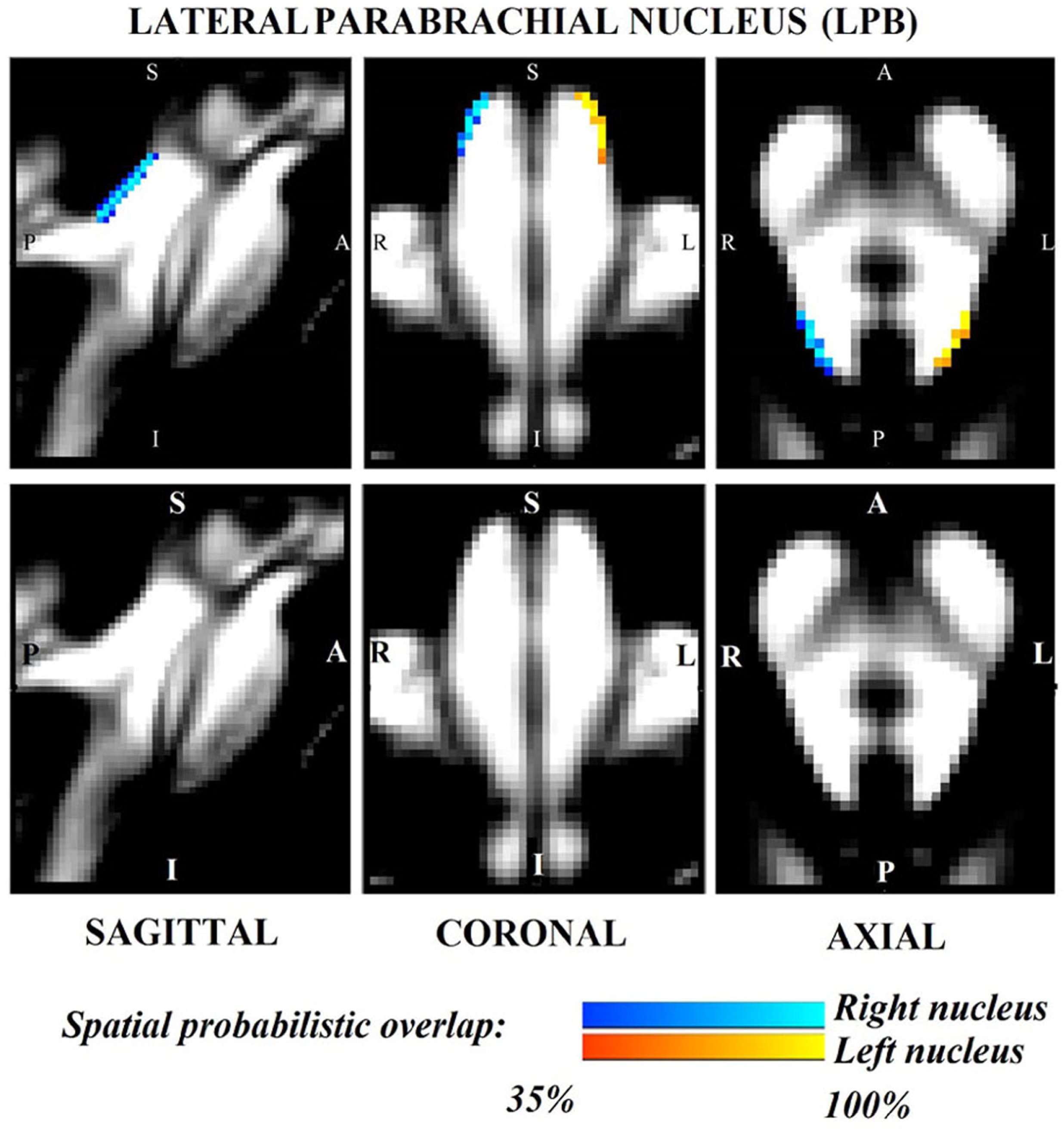
Probabilistic (n = 12) atlas label in MNI space of the lateral parabrachial nucleus. (right nucleus: blue-to-cyan; left nucleus: red-to-yellow). Very good (i.e. up to 100 %) spatial agreement of labels across subjects was observed indicating the feasibility of delineating the probabilistic label of these nuclei involved in arousal and autonomic functions.

**Figure 2.**
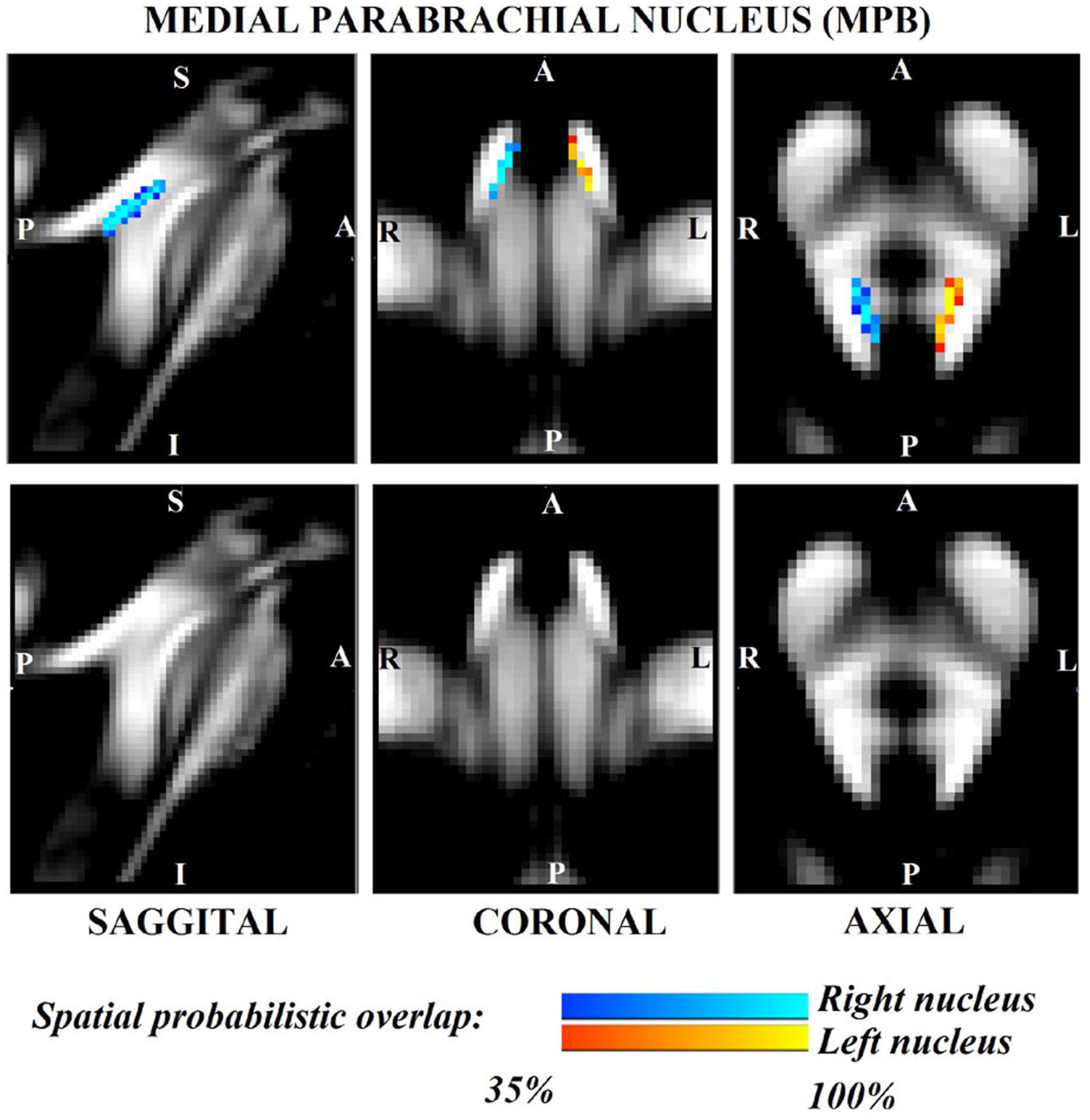
Probabilistic (n = 12) atlas label in MNI space of the medial parabrachial nucleus. (right nucleus: blue-to-cyan; left nucleus: red-to-yellow). Very good (i.e. up to 100 %) spatial agreement of labels across subjects was observed indicating the feasibility of delineating the probabilistic label of these nuclei involved in arousal and autonomic functions.

**Figure 3.**
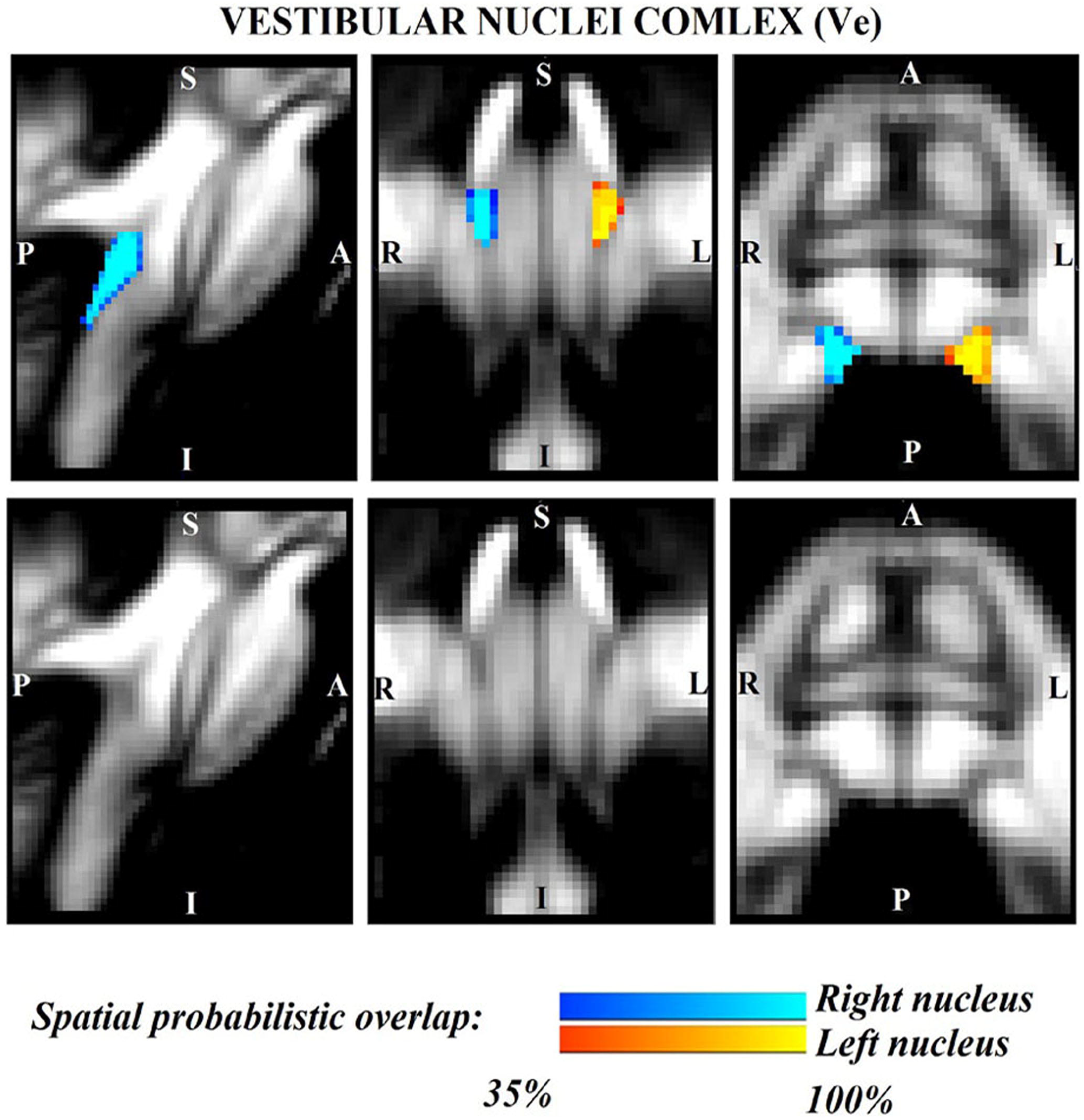
Probabilistic (n = 12) atlas label in MNI space of the vestibular-nuclei complex. (right nuclei complex: blue-to-cyan; left nuclei complex: red-to-yellow). Very good (i.e. up to 100 %) spatial agreement of labels across subjects was observed indicating the feasibility of delineating the probabilistic label of this complex of nuclei involved in vestibular functions (e.g. postural, oculo-motor control). Note that we did not have enough resolution/contrast to easily discriminate between *individual* (e.g. superior, lateral, medial, inferior) vestibular nuclei within this complex.

**Figure 4.**
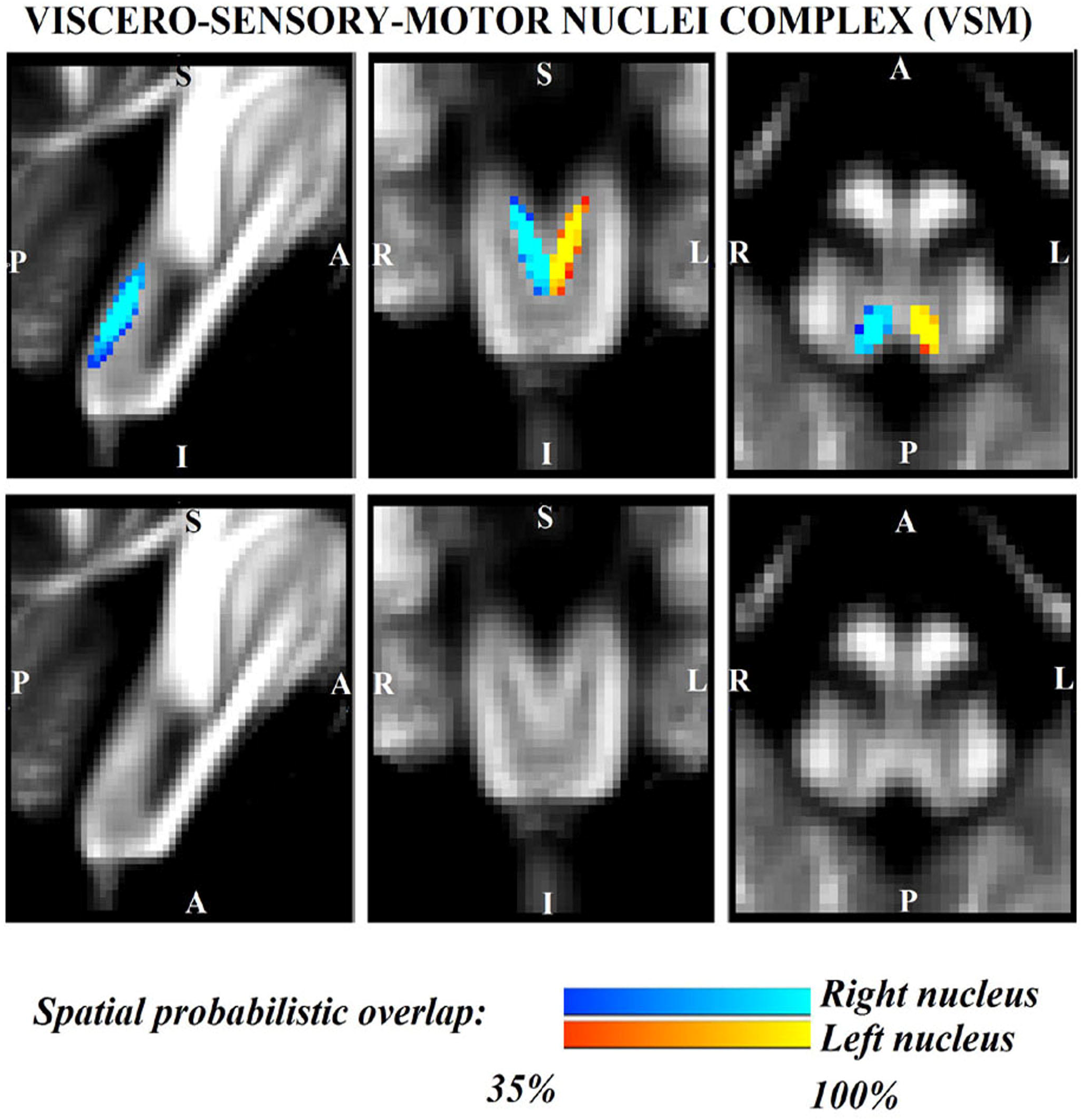
Probabilistic (n = 12) atlas label in MNI space of the viscero-sensory-motor-nuclei complex. (right nuclei complex: blue-to-cyan; left nuclei complex: red-to-yellow). Very good (i.e. up to 100 %) spatial agreement of labels across subjects was observed indicating the feasibility of delineating the probabilistic label of this complex of nuclei. Note that we did find good contrast showing V-shaped nuclei in the coronal section, which matched the exact description of nuclei from literature (Olszewski and Baxter 1954); yet, we did not have enough resolution/contrast to easily discriminate between *individual* nuclei within this complex (i.e. solitary nucleus, vagus nerve nucleus, hypoglossal nucleus, prepositus, intercalated nucleus, and interpositus).

For each nucleus, the average modified Hausdorff-distance assessing the inter-rater agreement and the internal consistency of nuclei atlas labels is shown in Figure 5. The modified Hausdorff-distance and the internal consistency were found to be below (p < 10^−7^) unpaired one-tailed t-test) the linear spatial imaging resolution (1.1 mm), thus validating the generated probabilistic nuclei labels.

**Figure 5.**
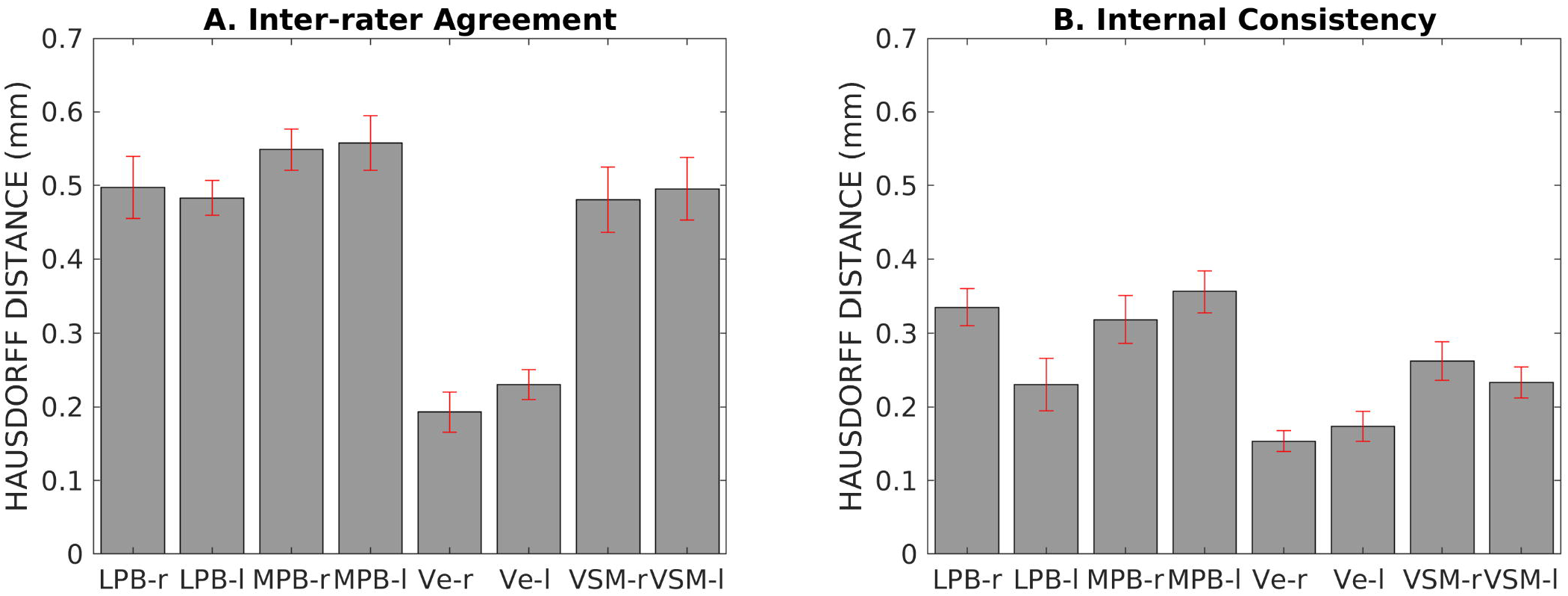
Atlas validation. We show: **A.** The inter-rater agreement of nuclei segmentations (bar/errorbar = mean/s.e. modified Hausdorff distance across 12 subjects); **B.** The internal consistency of nuclei labels across subjects (bar/errorbar = mean/s.e. modified Hausdorff distance across 12 subjects). The labels of the LPB-r/l, MPB-r/l, Ve-r/l and VSM-r/l displayed good spatial overlap across raters and subjects (the modified Hausdorff distance was smaller than the spatial imaging resolution), thus validating the probabilistic nuclei atlas.

The volume (mean ± s.e. across subjects) of each final label in native space and nuclei volumes from the literature (Paxinos et al. 2012) are shown in Table 1. The volume of MPB, Ve and VSM labels did not differ from Paxinos’ volumes (p-value < 0.05), whereas the volumes of LPB were larger than volumes computed from the Paxinos atlas (Paxinos et al. 2012). In Table 1, we report details of volume computation of LPB, MPB, Ve, VSM nuclei and of nuclei sub-regions, as obtained from the Paxinos atlas.

**Table 1.**
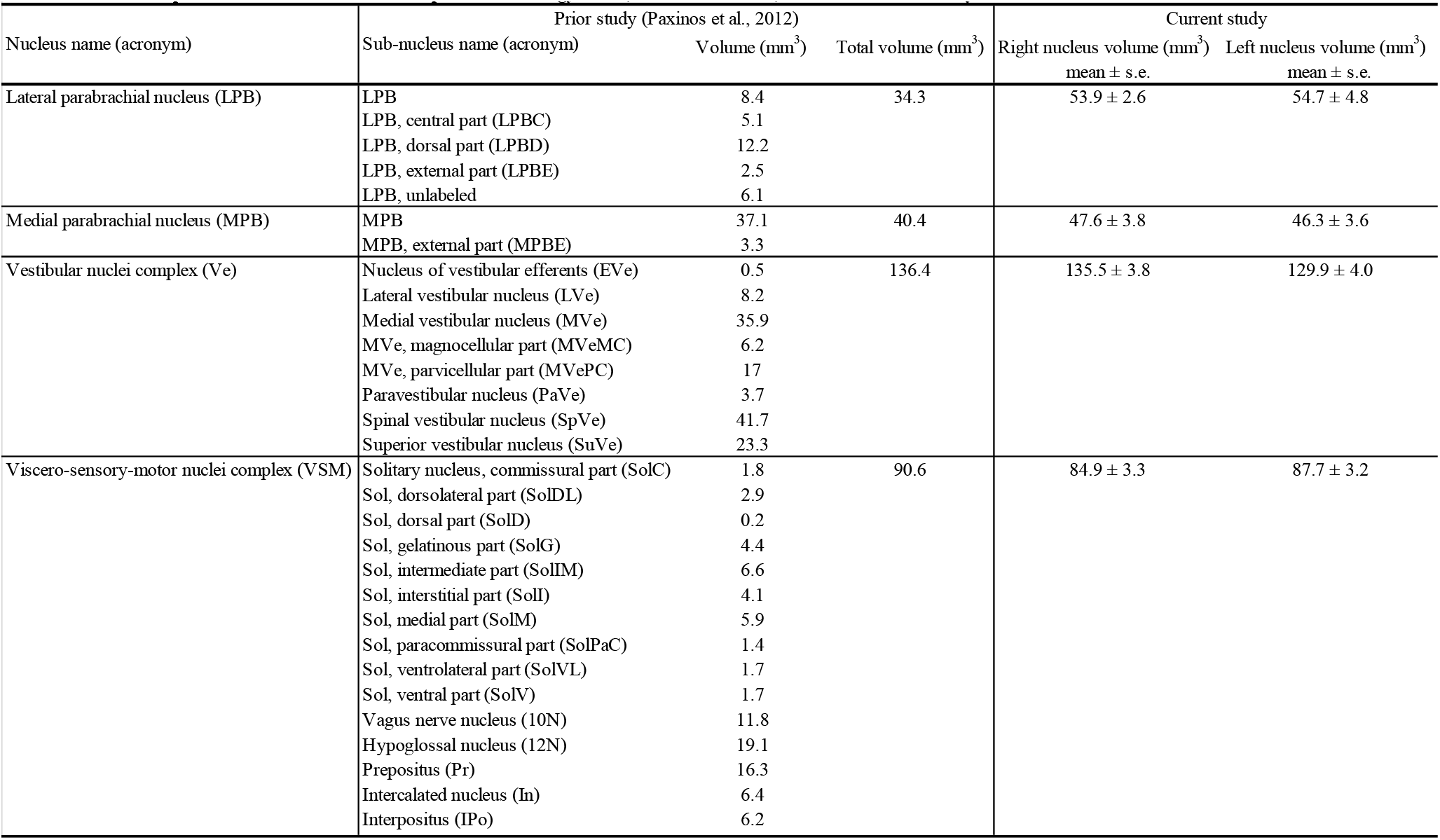
Detailed computation of nuclei volumes from previous histology atlas (Paxinos et al. 2012) and from current study

In Figure 6, we show a preliminary validation of the LPB and MPB *in vivo* nuclei label location and of their microstructural properties by the use of histology of a postmortem human brainstem specimen. Specifically, the FA map of MRI showed the hypointense region (as expected for gray matter nuclei) of LPB and MPB lining the hyperintense region of the superior cerebellar peduncle (SCP). Voxels of intermediate intensity in between SCP and parabrachial nuclei could be observed and could be attributed to partial volume effects. Nissl stain showed obvious difference between sparsely stained white matter of SCP and more densely stained grey matter of LPB and MPB displaying the distribution of neuronal cell bodies within these nuclei, further validating our nuclei localization. In the Gallyas staining, we found similar results where white matter of SCP could be demarcated from parabrachial nuclei based on their argyrophilic properties (Figure 6). These initial findings appear promising and need to be further extended for validation of Ve and VSM.

**Figure 6.**
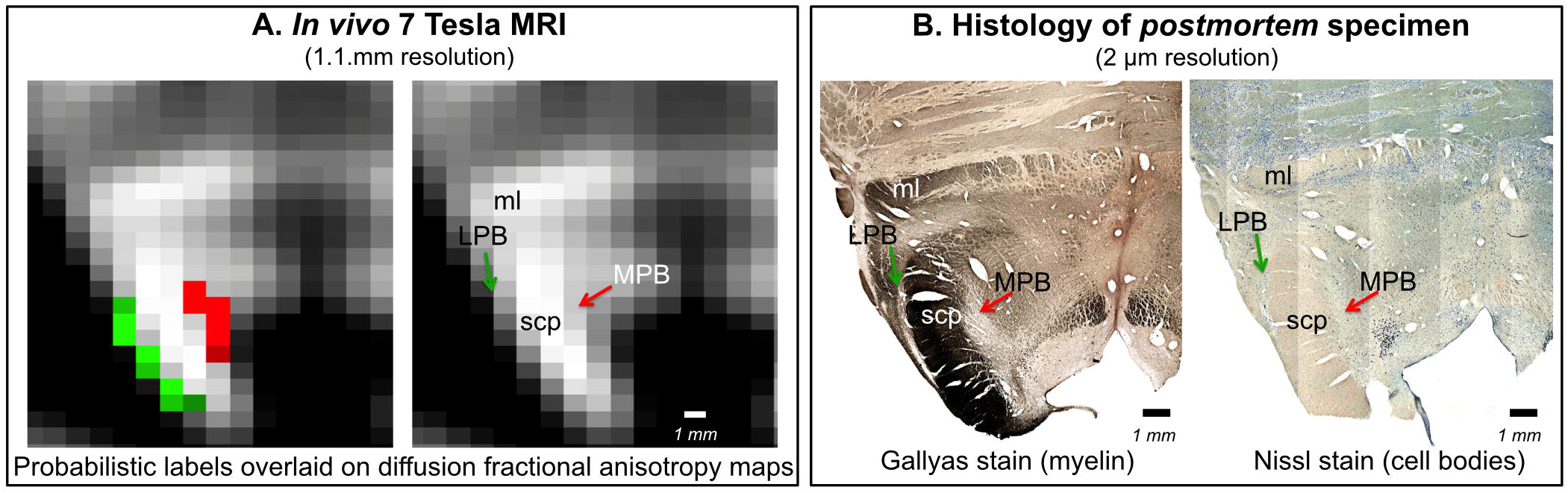
*In vivo* nuclei atlas validation by preliminary histology of postmortem brainstem specimens. We provide further validation for the LPB and MPB brainstem nuclei delineations. As expected for gray matter regions, LPB and MPB were areas of: **A.** hypointensity in *in vivo* 7 Tesla FA MRI maps; **B.** hypointensity in myelin stains (left) and hyperintensity in cell body stains (right) of a postmortem brainstem specimen. In each panel, the LPB and MPB bounded, respectively, laterally and medially the white-matter superior cerebellar peduncle (SCP; anterior to the SCP the medial lemniscus is labeled as ml).

## 4. Discussion

In the present work we demonstrated the feasibility of single-subject delineations of LPB, MPB, Ve and VSM nuclei in living humans using 7 Tesla MRI. Further, we created a probabilistic atlas of these nuclei after precise coregistration to stereotaxic neuroimaging space. Finally, we validated the generated atlas by assessing the inter-rater agreement, the consistency across subjects, and the volumes of the delineated labels. Our findings of most nuclei volumes match previously reported studies in literature, further strengthening our nuclei delineation. As a preliminary *ex-vivo* validation, we performed histological staining of a few nuclei (LPB, MPB), which showed remarkable consistency to *in-vivo* MRI. Here, we first discuss the strengths and limitations of nuclei delineations, then the nuclei function and the potential impact of the generated atlas.

### 4.1 Strengths and limitations of nuclei delineations

We delineated LPB, MPB, Ve and VSM nuclei based on high contrast FA maps and T_2_–weighted images obtained at 7 Tesla. As expected from postmortem atlases, LPB bounded laterally and superiorly (in its most caudal aspect) the superior cerebellar peduncle, medially the CSF and inferiorly the pedunculotegmental nucleus. The MPB laid medial to the superior cerebellar peduncle, superior to vestibular nuclei and lateral to a gray matter area containing the locus coeruleus, the laterodorsal tegmental nucleus and the central gray of the rhomboencephalon. The superior, medial, lateral and spinal vestibular nuclei were not clearly visible as individual nuclei, yet as a complex with homogeneous FA. The Ve nuclei complex, in line with Paxinos atlas, extended from the caudal tip of the MPB at the ponto-medullary junction, to the medulla at the level of the mid inferior olivary nucleus; it was medial to the inferior cerebellar peduncle and ventral to the forth ventricle. Similarly, Sol, 12N, 10N and smaller nuclei Pr, In, IPo were delineated within the VSM nuclei complex, not individually. We observed a peculiar diffusion FA contrast of the VSM, which displayed intermediate FA values between neighboring gray matter, such as the medullary reticular formation, and white matter, such as the medial longitudinal fasciculus. We speculate that the partial volume effect of the VSM with the solitary tract, nerves 10 and 12 might be at the origin of this contrast. In line with the Paxinos atlas (Paxinos et al. 2012), the VSM nuclei complex extended from the ponto-medullary junction, next to the Ve to the inferior medulla at a level of the lower tip of the inferior olivary nucleus, and was bounded dorsally by the CSF. Interestingly, on a coronal view, both VSM bilateral nuclei appeared as an inverted V-shape as described in other studies (Olszewski and Baxter 1954; Naidich et al. 2009; Paxinos et al. 2012). Future development using alternative contrasts and higher spatial resolution might enable the disambiguation of individual nuclei within both the Ve and VSM.

The quantitative validation using the inter-rater agreement and internal consistency of *in vivo* probabilistic labels provided positive results for all nuclei. Comparison of the probabilistic label volume with the volume derived from the Paxinos histology-based atlas drawings (Paxinos et al 2012) was quite good for MPB, Ve and VSM, yet there was some mismatch for LPB. We ascribe this discrepancy most probably to partial volume effects, because our LPB labels delineated using 1.1 mm isotropic resolution voxels might contain CSF and a stripe of the superior cerebellar peduncle adjacent to the LPB. The LPB is indeed a very thin stripe of gray matter with a width ranging between approximately 0.5 to 1 mm.

For proper applicability of this atlas, note that a precise coregistration to conventional MRI is needed. Nonetheless, coregistration of subcortical structure is less trivial than coregistration of cortical foldings (Fischl et al. 2008; Greve and Fischl 2009). Moreover, with the development of new registration tools (Klein et al. 2009; Avants et al. 2011) and of multi-modal atlases in MNI space (Varentsova, Zhang, and Arfanakis 2014) including diffusion-based contrast and T_2_-weighted contrast, beyond the original MNI T_1_-weighted contrast, we predict better accuracy of brainstem coregistration, for example from native single-subject space to MNI space. Same-modality coregistration of single-subject images to target stereotaxic images has provided promising outcomes for single-subject FA maps to the IIT FA map, as used here. Recent work by our group (Bianciardi et al. 2015, 2018), has demonstrated the feasibility of generating a template in MNI space and functional connectome of tiny brainstem structures, thus proving the feasibility of accurately coregistering these small nuclei across subjects to a common template.

### 4.2 On the nuclei function and the potential impact of the generated atlas

We delineated LPB and MPB nuclei located at the mesopontine junction and involved in chemoreception, nociception, stress, autonomic control, aversive behaviors and arousal functions (Spector 1995; Bester et al. 1997; Reilly and Trifunovic 2001; Gauriau and Bernard 2002; Chamberlin 2004; Pattinson et al. 2009; Fuller et al. 2011; Kaur et al. 2013; Davern 2014; Myers et al. 2017). For instance, functional studies (Pattinson et al. 2009; Zuperku et al. 2017) show the involvement of LPB and MPB nuclei in respiratory control; however, their roles could not be segregated due to unavailability of *in-vivo* human atlas specifically delineating these labels at high-resolution (Pattinson et al. 2009). MPB relays information from the taste area of the solitary nucleus to the ventral posteromedial nucleus of the thalamus and forebrain (Sclafani et al. 2001; Naidich et al. 2009), while the LPB relays viscero-sensory information. Owing to multi-functional involvement of sub-nuclei of LPB and its overlapping role with MPB, we speculate our atlas might help in refining definite roles for these nuclei in functional imaging in human subjects.

Brainstem vestibular and autonomic nuclei display an intricate wiring diagram with other brainstem nuclei and with the rest of the brain, as shown by animal and *ex vivo* work (Carey D. Balaban 2002; Carey D. Balaban, Jacob, and Furman 2011; Staab, Balaban, and Furman 2013). Alteration in the connectivity of these nuclei is the hallmark of several brain disorders (Millan 2002; Niblock et al. 2004; Brown, Lydic, and Schiff 2010; Linnman et al. 2012; Mello-Carpes and Izquierdo 2013; Satpute et al. 2013), including vestibular disorders and anxiety (C. D. Balaban and Thayer 2001; Carey D. Balaban 2004; Bächinger et al. 2019). According to the National Institutes of Health National Institute on Deafness and other Communication Disorders (NIDCD) (“2012-2016 NIDCD Strategic Plan” 2015), chronic vestibular disorders (including chronic imbalance and dizziness) affect about 5 % of the American adult population, their mechanisms are not fully understood (Kitahara et al. 1997; Ris et al. 1997; Godemann et al. 2005; Heinrichs et al. 2007; Best et al. 2009; Dutia 2010; E J Mahoney, Edelman, and D Cremer 2013; Cousins et al. 2014) and treatment with serotonergic antidepressants and vestibular habituation are only partially successful (Staab, Balaban, and Furman 2013). Adverse vestibular-autonomic interactions (Fischl et al. 2008; Staab et al. 2014; Indovina et al. 2014, 2015; Riccelli, Indovina, et al. 2017; Riccelli, Passamonti, et al. 2017a, 2017b; Nigro et al. 2018; Passamonti et al. 2018) appear to precipitate and perpetuate chronic vestibular disorders, crucially underlying the pathophysiologic process of these disorders. Our findings offer potential benefits to investigate the connectivity pathways of Ve and autonomic nuclei in living humans on widely available 3 Tesla scanners, and expand our knowledge of successful compensation for acute vestibular events versus development of chronic vestibular disorders.

Solitary nuclei integrate interoceptive and viscero-sensory input with descending affective and cognitive information from the limbic forebrain (Rinaman and Dzmura 2007). Early studies showed solitary nuclei role in autonomic control, however recent literature indicates their involvement in plethora of behavioral and neuroendocrine processes (Rinaman 2010), thereby further providing impetus for precise delineations of these nuclei in living humans. Solitary nuclei have been involved in behaviors relating to fear memory, anxiety and depression (Miyashita and Williams 2002; Ghosal et al. 2014) along with modulating behavioral responses to stress, which is also governed by parabrachial nuclei. Based on these growing evidences of overlapping functions and differential response of these nuclei to stimuli and their mode of action, we advocate that the present study might help with nuclei localization and their future functional assessment, as well as facilitate the study of various neurological disorders and neurosurgical planning.

### 4.3 Summary, conclusion and future directions

In summary, we foresee that the generated probabilistic atlas of LPB, MPB, Ve, and VSM in stereotaxic space —representative of younger human adults— might be a useful tool for improving localization of brainstem nuclei involved in autonomic, vestibular and viscero-sensory-motor functions. Researchers and clinicians will be able to shift from the difficult and imprecise task of extrapolating locations from *ex vivo* atlases to this new, user-friendly 3D, probabilistic, and deformable tool to identify the location of brainstem nuclei more precisely in *in vivo* images derived from conventional 3 Tesla MRI scanners. After its release on public repositories of neuroimaging data and tools, users will be able to precisely co-register the atlas onto single-subject 3 Tesla MRI, just as existing atlases (e.g., AAL, Harvard Oxford (Desikan et al. 2006; Destrieux et al. 2010) are now used. This will make the atlas accessible to researchers and clinicians who use widely available 3 Tesla scanners, and study brainstem mechanisms in health, vestibular and balance disorders, impairment in autonomic and viscero-sensory-motor function, sleep and anxiety disorders, as well as neurodegenerative disease.

## 5. Acknowledgements

This work was supported by the following sources of funding: National Institute on Deafness and other Communication Disorders R21 DC015888, National Institutes of Health (NIH) National Institute for Biomedical Imaging and Bioengineering K01 EB019474 and P41 EB015896, National Cancer Institute U01 CA193632, and the Massachusetts General Hospital Claflin Distinguished Scholar Award. This work was also in part made possible by the resources provided by Shared Instrumentation Grants 1S10RR023401, 1S10RR019307, and 1S10RR023043, Italian Ministry of Health (PE-2013-02355372).

